# scGCN: a Graph Convolutional Networks Algorithm for Knowledge Transfer in Single Cell Omics

**DOI:** 10.1101/2020.09.13.295535

**Authors:** Qianqian Song, Jing Su, Wei Zhang

## Abstract

Single-cell omics represent the fastest-growing genomics data type in the literature and the public genomics repositories. Leveraging the growing repository of labeled datasets and transferring labels from existing datasets to newly generated datasets will empower the exploration of the single-cell omics. The current label transfer methods have limited performance, largely due to the intrinsic heterogeneity and extrinsic differences between datasets. Here, we present a robust graph-based artificial intelligence model, single-cell Graph Convolutional Network (scGCN), to achieve effective knowledge transfer across disparate datasets. Benchmarked with other label transfer methods on totally 30 single cell omics datasets, scGCN has consistently demonstrated superior accuracy on leveraging cells from different tissues, platforms, and species, as well as cells profiled at different molecular layers. scGCN is implemented as an integrated workflow as a python software, which is available at https://github.com/QSong-github/scGCN.

## Introduction

Single-cell omics technologies are increasingly used in biomedical research to provide high resolution insight into the complex cellular ecosystem and underlying molecular interconnectedness [1-3]. Leading this new wave of omics is the single-cell RNA sequencing (scRNA-seq) that allows measurement of transcriptome in thousands of single cells from multiple biological samples under various conditions [4-8]. Other single-cell-based assays include Single-cell Assay for Transposase-Accessible Chromatin using sequencing (scATAC-seq) to profile cellular heterogeneity at the epigenetic level [9-11], which gains further insights into the elucidation of transcriptional regulators [12, 13]. These technological developments allow profiling multiple molecular layers at single-cell resolution and assaying cells from multiple samples under different conditions.

The fast advance of single-cell technologies leads to the rapid growth of single cell omics data. As more and more single-cell data becomes available, there is an urgent need to leverage existing data with the newly generated data in a reliable and reproducible way, learning from the established single-cell data with well-defined labels as reference, and transferring labels to new datasets to assign cell-level annotations [10, 11]. However, existing datasets and new datasets are often collected from different tissues and species [14, 15], under various experimental conditions, generated by different platforms [16, 17], and in the form of different omics types [18]. Thus a reliable and accurate knowledge transfer method must overcome the following challenges: 1) the unique technical issues of single-cell data (*e.g*., dropouts and dispersion) [19-22]; 2) batch effects arisen from different operators, experimental protocols [23], and technical variation (*e.g*., mRNA quality, pre-amplification efficiency, technical settings during data generation) [24-26]; 3) intrinsic biological variances associated with different tissues, species, and molecular layers such as RNA-seq and ATAC-seq.

To address these challenges in transferring labels across different datasets, several methods have been developed. The most commonly used are Seurat [27, 28] and the recently reported Conos [29], scmap [30], and CHETAH [31]. Seurat is a well-established, widely used toolkit for single cell genomics, which recently provides specialized approaches to transfer labels across substantially different single-cell samples [28]. Conos generates a joint graph representation by pairwise alignments of samples, to propagate labels from one sample to another. scmap learns cell types by measuring the maximum similarity between reference dataset with well-annotated cells and unknown datasets [32]. Guided by the reference data, CHETAH identifies a classification tree for a top-to-bottom classification in unannotated data. Whereas valuable in different settings, these methods exhibit limited capability and performance, partially due to that these methods only extract shared information from individual cells but ignore higher-order relations between cells. Such topological cell relation can be well captured by the Graph Convolutional Networks (GCN) [33]. Recently, GCN and its related methods have been successfully applied in single cell and disease [34-37], showing that inclusion of GCN enables to learn the high-order representation and topological relations of cells that improve performance.

Here, we present a novel, graph-based artificial intelligence model, termed single-cell Graph Convolutional Network (scGCN). We provide evidence that scGCN allows for reliable and reproducible integration of single-cell datasets and transferring labels across studies. Thus, knowledge learned from well-characterized datasets in previous studies can be transferred to and provides insights in new studies. Using a wide range of different single-cell omics datasets, including data from different tissues, species, sequencing platforms, and molecular layers (such as RNA-seq and ATAC-seq), we demonstrate that scGCN outperforms other methods in accuracy and reproducibility. In addition, we provide the implementation software of scGCN, which is compatible with various single-cell datasets for accurate cell type identification.

## Results

### Overview of the scGCN

Knowledge learned from existing single-cell datasets is often represented as cell labels. Examples of cell labels are cell type, developmental state, activation status, cellular functionality, and signaling pattern. The scGCN approach leverages well-characterized single-cell data as reference to infer such cellular-level knowledge in query dataset, *i.e*., label transfer, through semi-supervised learning (**Figure 1**). First, scGCN learns a sparse and hybrid graph of both inter- and intra-dataset cell mappings using mutual nearest neighbors of canonical correlation vectors that project different datasets onto a correlated low-dimensional space, which enables the identification and propagation of shared information between reference and query data. Subsequently, based on the constructed graph, semi-supervised GCN is used to project cells of both reference and new datasets onto the same latent space so that cells with the same labels present in the same population. Consequently, cell labels in query data are predicted and learned from reference data. Details are described in Methods.

**Figure 1.**
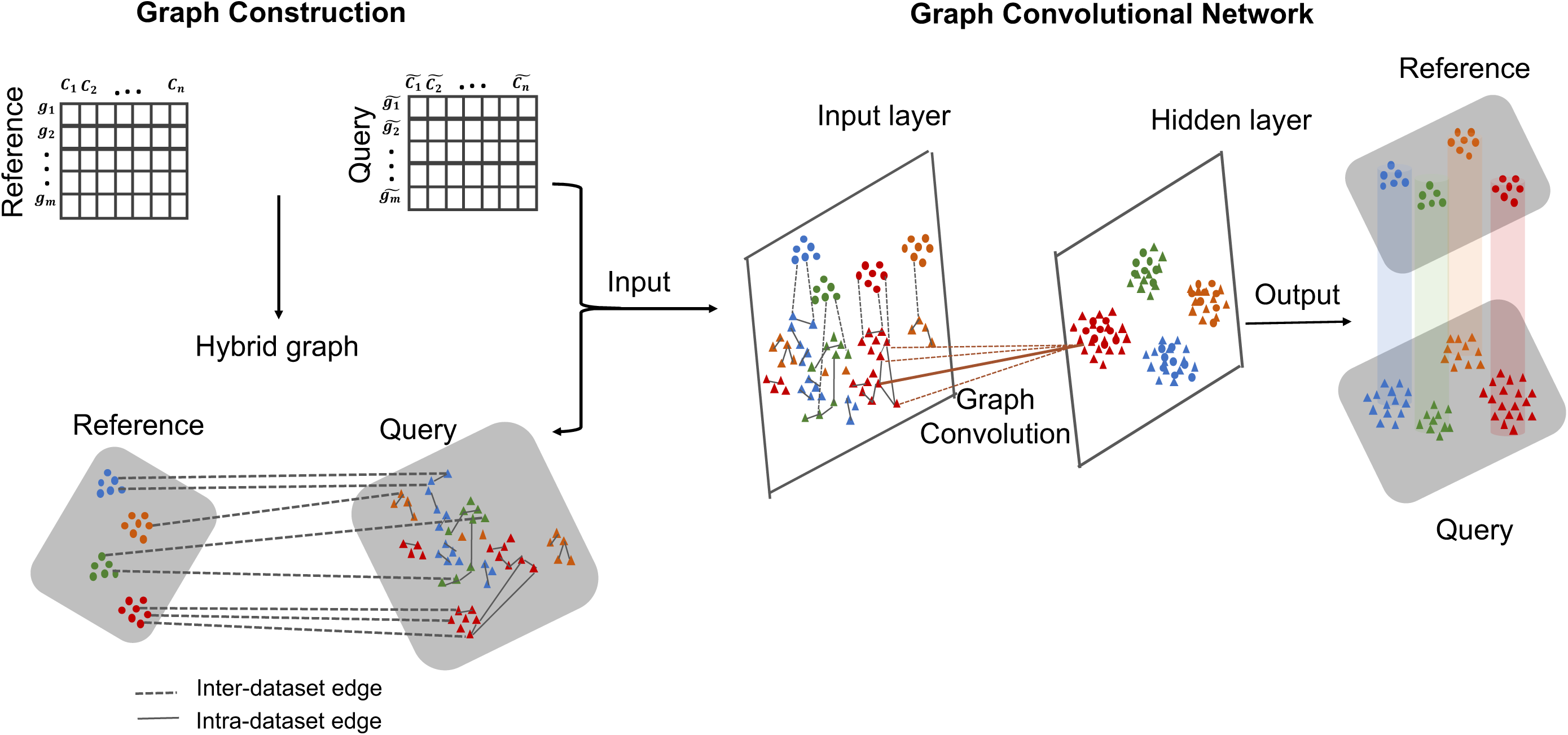
Schematic overview of scGCN for transferring labels from reference data to query data. scGCN first learns a hybrid graph of both inter-dataset (dash line) and intra-dataset (solid line) cell mappings. In the hybrid graph, round solid-colored dots represent cells of reference data, while the triangle shapes represent cells from query dataset. Based on the hybrid graph, semi-supervised GCN is used to project cells of both reference and query datasets onto hidden layer so that cells with the same labels present in the same subpopulation. Thus, cell labels in query data are predicted and learned from reference data.

### Performance of label transfer within datasets

To evaluate the performance of scGCN, we benchmark it against other methods, including Seurat [28], Conos [29], scmap [30], and CHETAH [31]. For quantitative benchmarks, we first use 10 scRNA-seq datasets that vary in cell numbers, tissues, species, and sequencing technologies. These datasets represent different scenarios and challenges in cell label transfer. For each dataset, we randomly select 50% of its cells as the reference data and apply the above methods to learn the labels of the other 50% of cells as the query data.

We evaluate the performance of each method using the accuracy score (Acc), which is defined as the proportion of correctly predicted cells among all cells in the query data. Our results show that scGCN consistently demonstrates better performance than other methods across datasets and achieves the highest accuracy (mean Acc = 91%, **Figure 2A**), which is significantly higher than other methods (Seurat: 87%; Conos: 83%; scmap: 82%; CHETAH: 77%) (*P* value = 0.0019, 0.0058, 0.0039, and 0.002, respectively). scGCN demonstrates particularly higher accuracy relative to these methods in some datasets. For example, Seurat shows a relatively lower accuracy score in GSE99254 dataset (Seurat: 77%; scGCN: 85%), while Conos performs poorly on the GSE108989 dataset (Conos: 53%; scGCN: 86%), and CHETAH performs poorly on the SRP073767 dataset (CHETAH: 62%; scGCN: 90%). Therefore, scGCN achieves the best performance in transferring labels accurately in the 10 benchmarking datasets.

**Figure 2.**
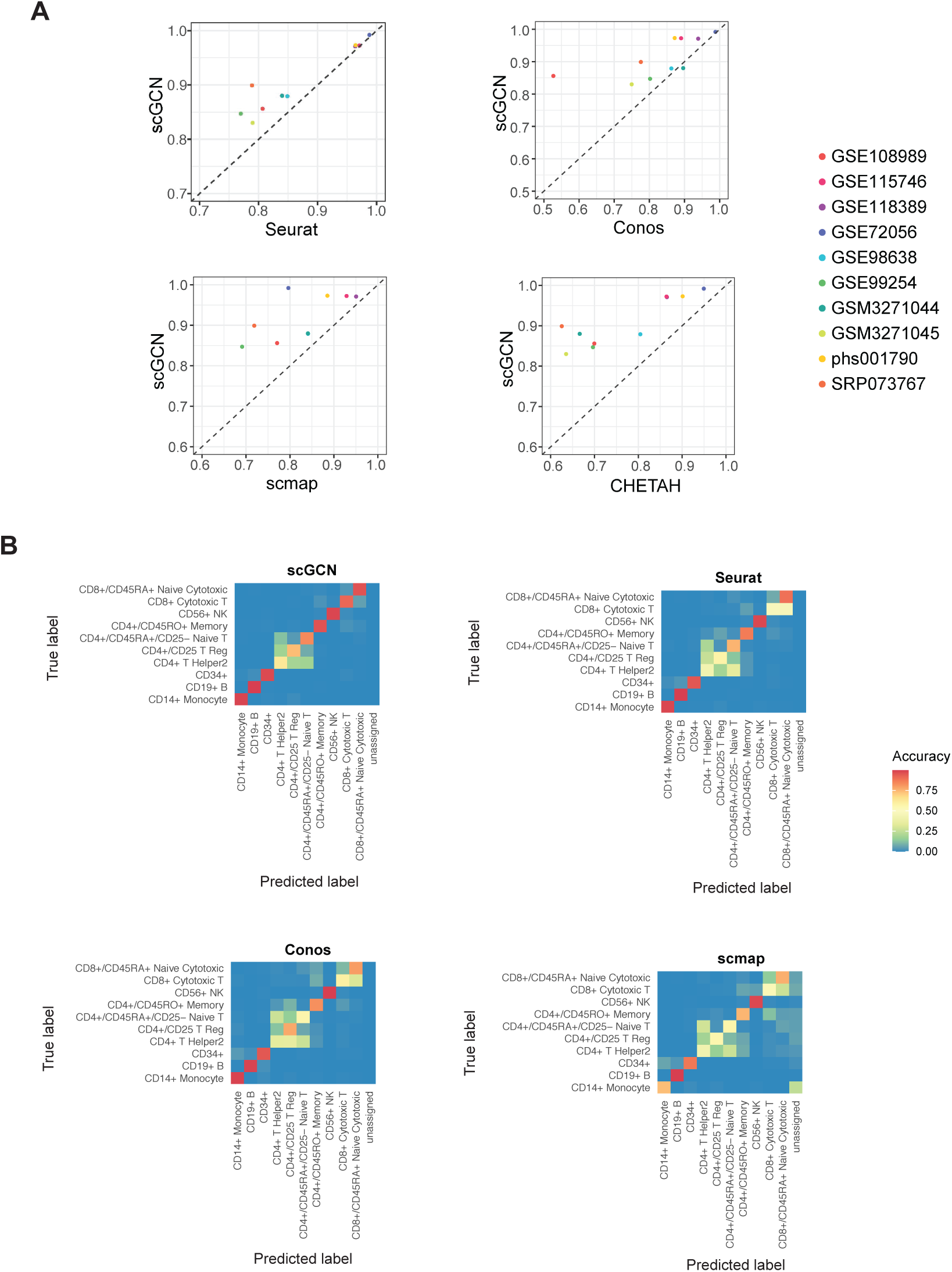
Performance of label transfer within datasets. **A**. Performance of scGCN and other methods (Seurat, Conos, scmap, and CHETAH) are measured by the accuracy score on 10 datasets. Each point represents the accuracy scores of scGCN versus an alternative method on one dataset. The dashed line represents equivalent accuracy between two methods. Dots above the dashed line represent that scGCN outperforms the corresponding method on these datasets. **B**. Heatmap of the accuracy matrix of each cell type, including different T-cell subtypes, B-cells, and monocytes, which are identified by different methods based on the single cell dataset (SRP073767).

To highlight the comparison regarding specific cell types, we use the SRP073767 dataset as an example, which has two subtypes of T cells (CD4+ T helper2 cells, CD4+/CD25 Treg cells) that every benchmarked method cannot distinguish accurately whereas scGCN relatively performs better (**Figure 2B, S1A**). In addition, Seurat and Conos cannot discriminate between CD4+ T helper2 cells and CD4+/CD45RA+/CD25− Naive T cells, nor the CD8+/CD45RA+ Naive Cytotoxic cells and CD8+ Cytotoxic T cells. scmap and CHETAH also inaccurately assigns CD4+ to CD8+ T cells, in addition to the low accuracy of monocytes classification and the unassigned cells. scGCN, in contrast, performs better than other methods in discerning these similar cell types.

### Transfer labels across datasets of different platforms

As emerging single cell data are generated by different experimental platforms, we test whether scGCN can be used to accurately transfer labels across datasets from different platforms. Here we include 12 paired reference-query datasets. Each pair of reference-query datasets are profiled using different scRNA-seq technologies.

Similarly, we use the accuracy score to evaluate the performance of each method. Based on the 12 reference-query datasets, the accuracy score of scGCN (mean Acc = 87%, **Figure 3A**) is consistently higher than Seurat (mean Acc = 82.2%) and Conos (mean Acc = 82.3%), and also significantly better than scmap (mean Acc = 66%; *P* value = 0.001433) and CHETAH (mean Acc = 58%, *P* value = 2.219e-05). Specifically, when querying the PBMC Smart-seq2 data from the reference of PBMC Dropseq data, both Seurat (Acc = 68%) and scmap (Acc = 60%) show relatively lower accuracy scores than scGCN (Acc = 77%). Similarly, Conos (Acc = 71%) and CHETAH (Acc = 32%) present lower accuracy than scGCN (Acc = 76%) when mapping MCA Smart-seq2 from MCA 10X data. Notably, when annotating the PBMC 10X V3 data using the PBMC Cel-seq data as reference, scGCN is the most distinctively superior than other methods. Although these two datasets are generated using different platforms, Uniform Manifold Approximation and Projection (UMAP) [38] reveals highly consistent transferred labels by scGCN for the cell populations of the PBMC 10X V3 query data (**Figure 3B**). Sankey diagram shows that the reference data is much smaller than the query data (**Figure 3C**), suggesting the effectiveness of scGCN even with small reference data.

**Figure 3.**
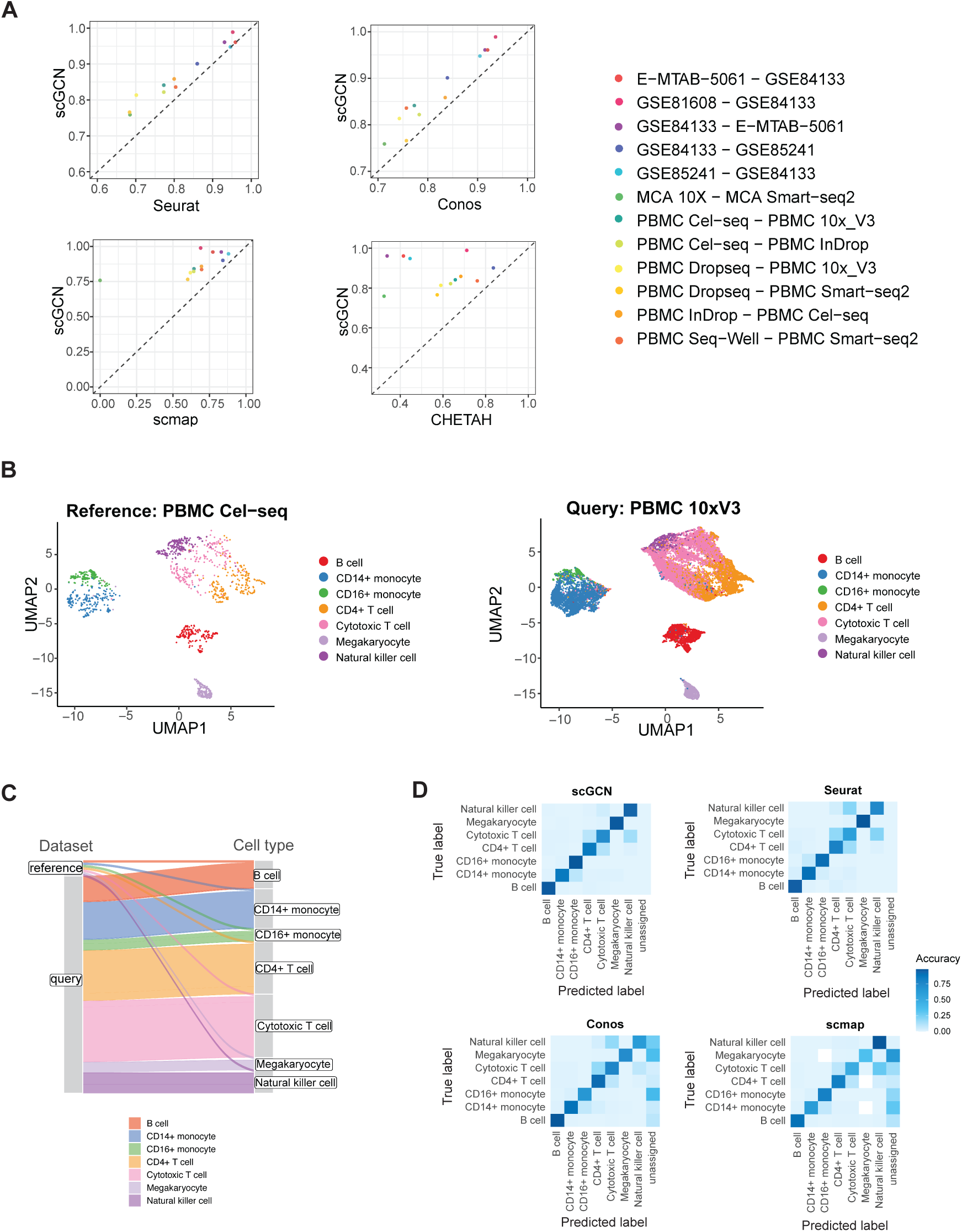
Performance of scGCN and its comparison with other methods using 12 cross-platform datasets. **A**. Performance of scGCN and alternative methods are measured by the accuracy score for 12 paired datasets. Each pair of reference-query datasets are profiled using different single-cell platforms. Each point represents the accuracy score for the query data of each paired datasets. **B**. scGCN produces UMAP projection of cells from Cel-seq as reference to annotate cells from 10x V2 as query. **C**. The height of linage line in the Sankey diagram reflects the cell number of each cell type in the reference data and query data. **D**. Heatmap shows the accuracy matrix of each cell type identified by different methods based on the Cel-seq data as reference and the 10x V2 data as query set.

We next use heatmap to present the accuracy of each cell type including B cells, CD4+ T cells, and cytotoxic T cells that are identified by different methods (**Figure 3D, S1B**). Seurat and CHETAH incorrectly assign most CD4+ T cells and natural killer cells to cytotoxic T cells. Some Megakaryocytes and CD16+ monocytes are assigned to other cell types by Conos and scmap. The closely related cell types such as natural killer cells and T cells, as well as CD14+ monocytes and CD16+ monocytes, are correctly discriminated by scGCN.

### Transfer labels across datasets of different species

We next evaluate the performance of scGCN across datasets from different species. We apply all benchmarking methods to 4 paired reference-query datasets. For each paired datasets, one consists of cells from mouse and the other from human tissues.

We first apply all five methods to identify the labels of query data in the 4 pairs of datasets. Then we compare the visualization of cells using the aggregated reference-query data by different methods (**Figure 4A**). Because CHETAH does not provide the aggregated data, we omit this method in the following comparisons of cell visualization. Due to the inherent noise and batch effects in the raw data, cells are not separated well, particularly for dataset 3, *i.e*. phs001790 (mouse) – GSE115746 (human), and dataset 4, *i.e*. GSE115746 (mouse) − phs001790 (human). Similarly, in the UMAP projections of Seurat and Conos, cells are not distinguished explicitly in datasets 3 and 4. scmap shows far less discernable results in these paired datasets (**Figure S1C**). In contrast, when using the aggregated data generated by scGCN, cell subpopulations are clearly discerned in the UMAP projections for all 4 scenarios.

**Figure 4.**
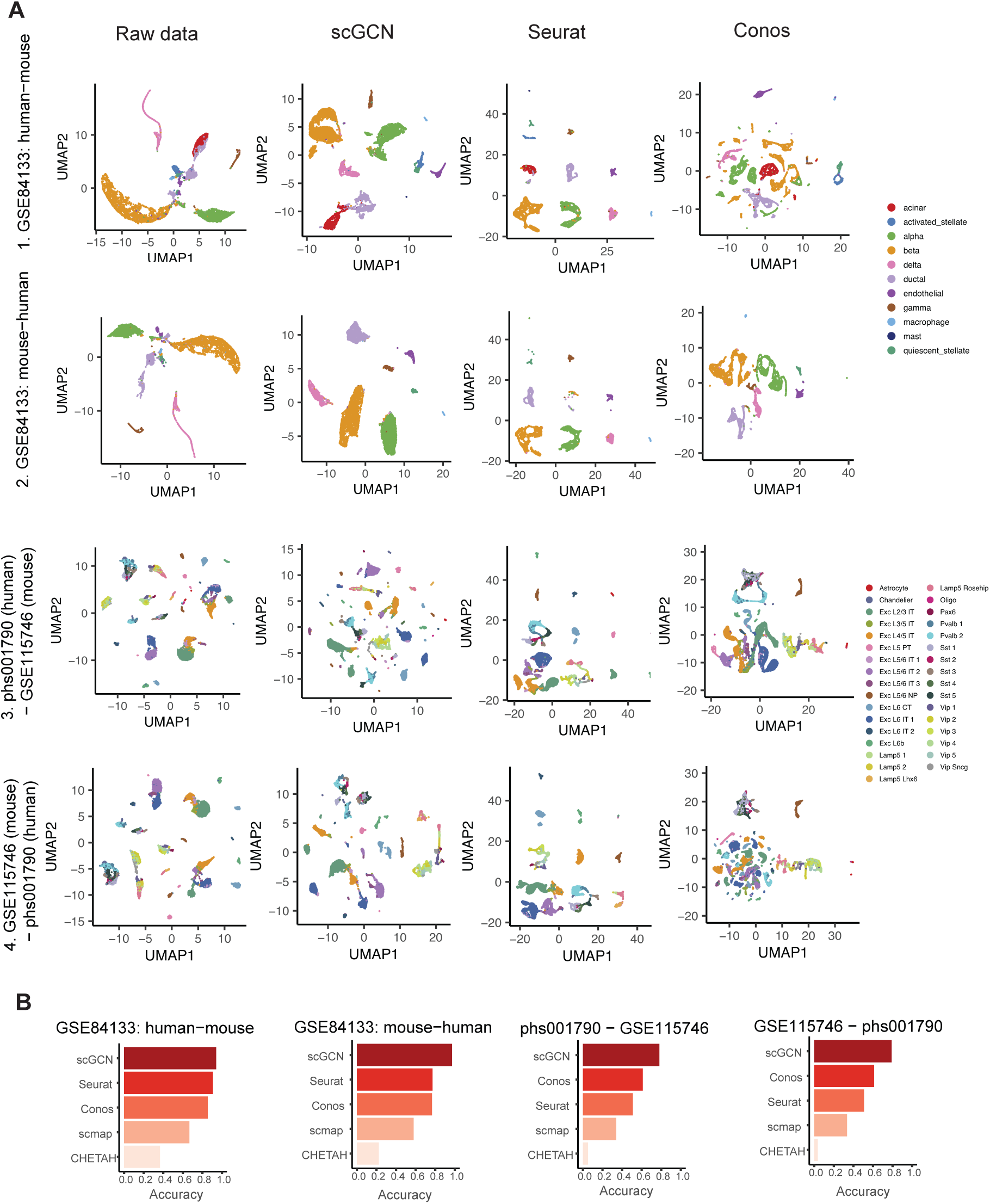
Performance of scGCN and its comparison with other methods using four cross-species datasets. **A**. UMAP projection of four paired cross-species datasets, based on the aggregated data by different methods. The top two rows represent the aggregated data using the human and mouse pancreas dataset GSE84133. First row: human data as the reference and mouse data as the query data. Second row: mouse data as reference and human data as query. Totally 11 cell type labels are visualized. The bottom two rows represent cells collected from human (phs001790) and mouse cortex (GSE115746), with human and mouse samples as the reference in the third and fourth rows, respectively. Totally 33 cell type labels are visualized. **B**. The bar plots show the performance of scGCN versus other methods (Seurat, Conos, scmap, and CHETAH) that are measured by the accuracy score for 4 paired cross-species datasets shown in **A**. Each bar represents the accuracy score of each method.

The UMAP projections suggest that the aggregated cells are aligned better by scGCN than the other methods. This observation is further confirmed by the accuracy score (**Figure 4B**). Specifically, scGCN shows the best performance with higher average accuracy (87%), compared with Conos (71%), Seurat (67%), scmap (48%), and CHETAH (20%). We find that Conos shows higher accuracy than Seurat in datasets 1 and 2, but lower in datasets 3 and 4. Quantitatively, in the 4 paired datasets, scmap and CHETAH produce less accurate results than those of Seurat and Conos. Together, these results show that scGCN performs consistently better in transferring labels (*e.g*. Seurat and Conos) across different species.

### Transfer labels across datasets of different types of omics

We next examine how well different algorithms transfer labels across different types of omics. Here we apply scGCN, Seurat, and Conos, but omit scmap and CHETAH as they are only designed for scRNA-seq data. Four open accessible paired datasets (A549, brain, kidney, and lung tissues) with scRNA-seq data as reference and scATAC-seq data as query are included for comparison. We use two evaluation metrics including the batch mixing entropy and the silhouette coefficient. The batch mixing entropy shows the mixing level of cells in the aggregated profiles of reference and query data [39]. A higher entropy value means better intermingling of cells from different batches, wherein the scRNA-seq and scATAC-seq data are regarded as two batches. The silhouette coefficient quantifies the separation of different cell types by calculating the silhouette widths of each cell type [39, 40]. A higher silhouette coefficient represents more differences between cell types while fewer variances within each cell type.

Using the above two metrics, we evaluate the transferred labels in each query dataset, *i.e*. scATAC-seq data. Compared to the other methods, scGCN shows better performance with higher mixing entropy (**Figure 5A**). Specifically, Seurat has higher mixing entropy than Conos in A549 data, but lower than Conos in the other three datasets. In the brain dataset, Conos shows comparable mixing entropy with scGCN. We then calculate the silhouette coefficients for all three methods (**Figure 5B**). In comparison with Seurat and Conos, scGCN has significantly and consistently higher silhouette coefficients. In most datasets, Seurat shows slightly higher silhouette coefficients than Conos.

**Figure 5.**
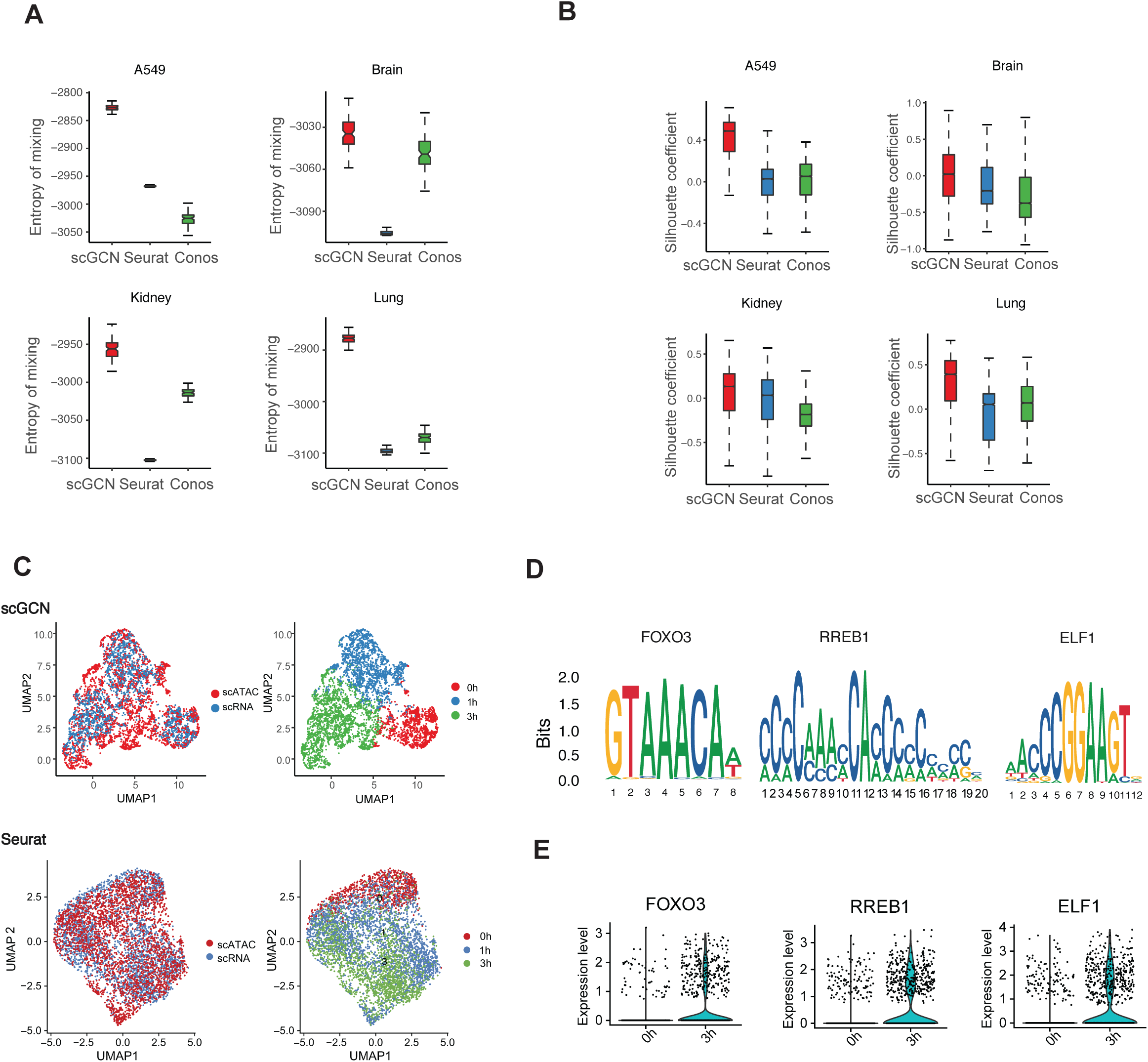
Performance of scGCN and its comparison with other methods using four paired cross-omics datasets. **A-B**. For each method, performance metrics including two indexes, i.e. batch effects entropy and silhouette index, are used to evaluate the level of cell mixing across datasets and the preservation of local structure within dataset. **C**. UMAP plots of A549 cells colored by dataset (scRNAseq, scATAC-seq) and by cell states (0h, 1h, 3h), after integration by scGCN and Seurat. **D**. Overrepresented DNA motifs are identified in 3h-specific accessibility peaks, with FOXO3, RREB1, and ELF1 motifs as the most highly enriched motifs. **E**. These motifs also exhibit upregulated expression in scRNA-seq cells at 3h.

To further visualize the joint alignment of scRNA-seq data and scATAC-seq data, we show the UMAP projection of the aggregated A549 cells with labels learned by different methods (**Figure 5C**). The cell subpopulations (0h, 1h, and 3h cells after 100 nM dexamethasone (DEX) treatment) are clearly discerned in the UMAP projection when using the aggregated data generated by scGCN. In contrast, Seurat and Conos are not able to explicitly distinguish different cell subpopulations (**Figure 5C, S2A**).

To characterize the differential accessible loci and uncover the transcriptional regulatory mechanisms in the scATAC-seq data of A549 cells, we perform the motif enrichment analysis to discover the cis-regulatory DNA sequences that differentially regulate the 3h cells after DEX treatment. Notably, comparing the 3h cells after DEX treatment with 0h cells, the specific loci of 3h cells is enriched with the binding motifs of FOXO3, REBB1, and ELF1 (**Figure 5D**). Moreover, we find that the transcription factor FOXO3 is upregulated in 3h cells (**Figure 5E**), which is a validated regulator that drives cell progression under the DEX treatment [41, 42]. KRT7 and WDR60 also show higher expression in 3h cells, which has been revealed *in vivo* of DEX treatment [43, 44] (**Figure S2B**). RREB1 and ELF1 exhibit upregulation in 3h cells compared to 0h cells (**Figure 5E**). These results suggest the potential roles of these identified transcriptional factors in maintaining and establishing the chromatin accessibility to express functional genes after treatment. Similar motif analysis is performed in mouse brain dataset, based on the integrated scRNA-seq and scATAC-seq datasets from the 10x Genomics Chromium system (**Figure S3**). In this data, overrepresented DNA motifs are identified in L4-specific accessibility peaks, with Foxp1, Egr3, and Smad3 motifs as the highly enriched motifs (**Figure S3A**), which also exhibit upregulated expression in L4 cell subtype (**Figure S3B**). Altogether, these results suggest that scGCN outperforms when transfer labels between single cell transcriptomics and epigenomics data.

## Discussion

Single-cell omics technologies have allowed biologists to gain unprecedented views into the individual cellular components of complex biological ecosystems [45, 46]. Facing the explosive and fast-growing single-cell data, there is a critical need to leverage the existing, well-characterized datasets as reference to ensure reliable and consistent annotations of new data. In this study, we report a graph-based artificial intelligence model, single-cell Graph Convolutional Network (scGCN) that empowers single cell omics researchers to use reference single-cell omics data to annotate new data through robust knowledge transfer approach. We provide evidence that scGCN allows for reliable and reproducible cell type transferring across datasets from different tissues, species, sequencing platforms, and molecular layers (such as RNA-seq and ATAC-seq). The scGCN software, which is publicly accessible, is compatible with various single-cell datasets for accurate label transfer.

It is recognized that apart from the technical aspects of computational analysis, the accuracy, robustness, and sensitivity of label transfer also rely on the quality of the reference datasets. Reference datasets used in this study are well-characterized in literature, covering various types of samples and application scenarios in biomedical research. Whereas the current study does not address the quality issues of the reference datasets, it is well accepted that accumulating data in the field will lead to development and definition of higher quality and experimental evidence-based reference datasets. Additionally, when annotating the PBMC 10X V3 data, scGCN reveals accurate cell mapping even when the reference data has small cell number, suggesting that scGCN is less insensitive to cell number than other methods, which can be further validated with more datasets for a systematic and stringent analysis.

Four commonly used algorithms, Seurat, Conos, scmap, and CHETAH, also possess the knowledge transfer functionalities. We compare the performance of scGCN with these algorithms and show that scGCN consistently outperform. From a technical perspective, scGCN provides some major advantages. First, different from the other methods, scGCN simultaneously utilizes features, graphical structures, and reference labels to overcome the batch effects, protocol differences, and other intrinsic differences (e.g. different species and omics types) among datasets. Second, scGCN transfers labels from labeled samples to unlabeled samples in a semi-supervised manner, which is more desirable than other unsupervised methods. Third, in every layer, scGCN nonlinearly propagates feature information from neighboring cells in the hybrid graph, which learns the topological cell relations and improves the performance of transferring labels by considering higher-order relations between cells.

In addition to the successful results, there are several aspects in which scGCN can be improved. First, as an artificial intelligence (AI) model, scGCN shows not only the merits of its kind, but also some limitations including the black-box nature of AI models [47-49], which can be addressed through downstream analysis such as differential gene identification and enrichment analysis that can ameliorate some of the problems and bring insights into the labeled cells. Second, as a graph model, improving the graph construction can further boost the model performance. Our graph construction based on mutual nearest neighbors reflects the state-of-art in single cell graph representation. As a fast-growing research field, new approaches of graph construction are emerging, which we will test and adapt in future versions of scGCN.

## Methods

### Data preprocessing

For each input data, we denote the dataset with known cell labels as reference data *X*_*R*_ ∈ 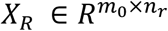 and the dataset that needs to be annotated as query data *X*_*Q*_ ∈ 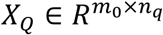, where *m*_0_ is the number of common gene features shared by *X*_*R*_ and *X*_*Q*_, *n*_*r*_ and *n*_*q*_ are the number of cells in reference data *X*_*R*_ and query data *X*_*Q*_. First, we identify the gene features that exhibit the most variability across different cell types in the reference data, which can be represented as heterogenous and prioritized features. As there are multiple cell types present in the reference data (the number of different cell types is annotated by *F*), we perform multi-class differential expression analysis to 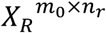 using Analysis of Variance (ANOVA) to identify the most variable genes across different cell types. Bonferroni correction is used to select the top *m* = 2, 000 genes in reference data with significant adjusted *P* value. In scGCN, all input data filters out non-variable gene features, thus the reference data and the query data become *X*_*R*_ ∈ 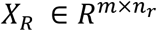 and *X*_*Q*_ ∈ 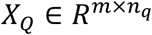.

### Graph construction

We use the mutual nearest neighbor (MNN) [50] approach and canonical correlation analysis [51] to construct a hybrid graph capturing the topological characteristics of single cells in both reference and query datasets. First, the reference dataset as well as the query dataset are standardized respectively. Then a reference-to-query graph (inter-dataset) as well as an internal query graph (intra-dataset) are constructed. The final hybrid graph is composed of these two graphs. The hybrid graph is then used as input for the scGCN model.

#### Standardization transformation

We first perform the standardized transformation for reference data, *i.e*.

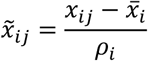

where *x*_*ij*_ ∈ *X*_*R*_ is the raw value of input reference data, 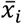 is the mean of *x*_*ij*_, and *ρ*_*i*_ is the standard deviation of *x*_*ij*_. Thus 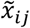 is the standardized value of feature *i* and cell *j*, where *i* ∈ {1,2, …, *m*} and *j* ∈ {1,2, …, *n*_*r*_}. After standardization, reference data *X*_*R*_ is represented as 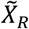. In the same way, raw data in *X*_*Q*_ is processed in the same manner that becomes 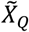.

#### Construction of the reference-to-query graph

For the scGCN method, one critical step is to construct an effective graph, which is represented as an adjacent matrix that best leverages the reference data and query data. Here we use the concept of mutual near neighbor (MNN) [50] and search for the MNNs from both reference and query data after simultaneous dimensionality reduction of reference and query datasets through canonical correlation analysis.

The goal of canonical correlation analysis is to simultaneously project the high-dimensional reference and query data into the same low dimensional space through two dataset-specific linear transformations so that the molecular patterns in both reference and query datasets that share the same biological meaning can be captured and represented uniformly in the low dimensional space. As 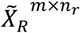 represents the standardized reference data and 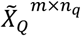 represents the query data. In order to project these two matrixes to a *k*-dimensional space where *k ≤ m*, we need to identify *k* pair of canonical correlation vectors *μ*_*i*_ of *n*_*r*_ dimension and *v*_*i*_ of *n*_*q*_ dimension, where *i* = 1, …, *k*, to maximize

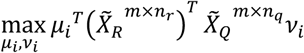

s.t. 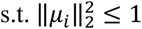 and 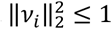. We use singular value decomposition (SVD) to calculate the *k* canonical correlation vector pairs which are associated with the *k* largest eigenvalues. Each pair of canonical correlation vector *μi* and *v*_*i*_ thus can be used to project the original data 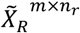 and 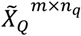 to the *i*’th dimension in the *k*-dimensional space, respectively. In this study we set *k* = 20.

The reference-to-query graph *A*^*RQ*^ is the constructed using MNN approach using the projected reference and query data. For a cell *i* in the reference data and another cell *j* in the query data, 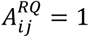 if and only if cell *i* belongs to the nearest neighbors of cell *j* and at the same time cell *j* is also one of the nearest neighbors of cell *i*. Otherwise 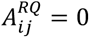. Thus, we construct the inter-dataset cell mapping between reference data and query data.

#### Construction of the query internal graph

Similarly, we also construct the internal graph *A*^*QQ*^ for cells in the query dataset using MNN approach and canonical correlation analysis with *k* = 20.

#### Construction of the hybrid graph

The final the hybrid graph 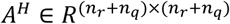 is then constructed by combining the reference-to-query graph (*A*^*RQ*^) and the query internal graph (*A*^*QQ*^). The hybrid graph *A*^*H*^ is used as the input adjacent matrix for scGCN model.

### scGCN method

We utilize the Graph Convolutional Network (GCN) [33] for semi-supervised learning and transferring labels from reference data to query data. Each cell is viewed as a node. The annotations of cell types in the reference data are the known labels, the goal of GCN is to predict the cell type annotations of the query data by using not only the features of each cell but also the information leveraging reference and query data, which are characterized as the above hybrid graph *A*^*H*^. Explicitly, the GCN model takes two inputs. One input is the hybrid graph structure learned above, which is represented as the adjacent matrix *A*^*H*^ ∈ *R*^*N×N*^ (see the **Graph construction)**. The other is the feature matrix *X* ∈ *R*^*m×N*^, where *N* = *n*_*r*_ + *n*_*q*_ is the total number of cells and *m* is the number of variable features selected in preprocessing. If the reference data is denoted as *X*_*R*_ ∈ 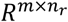 and the query data as 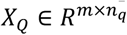, the input data matrix is extended to

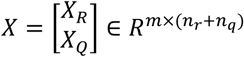

Herein, both features of reference and query data, as well as the information leveraging reference and query data are utilized in our model. With these two inputs, the GCN model is constructed with multiple convolutional layers.

For efficient training of GCN introduced in [33], the adjacent matrix *A*^*H*^ is modified as:

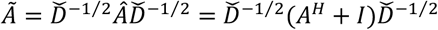

where *I* is the identity matrix, *Â* = *A*^*H*^ + *I*, and *Ď* is the diagonal degree matrix of *Â*. Specifically, each layer is defined as:

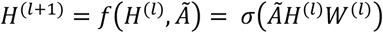

where *H*^(*l*)^ is the input and *W*^(*l*)^ is the weight matrix of the *l*-th layer, *σ*(·) is the non-linear activation function, and the input layer *H*^(0)^ = *X*. The labels of cells in the reference data are represented as an class indicator *y*_*lf*_ for cell *l* ∈ *𝒴*_*L*_ and label *f* ∈ {1, …, *F*}, where *𝒴*_*L*_ represents nodes with known labels, *F* represents the total number of different labels, and *y*_*lf*_ = 1 indicates cell *l* has the *f*’th label, and 0 as not of this label.

Specifically, for a three-layer GCN with *F* distinct labels, the forward propagation is realized as:

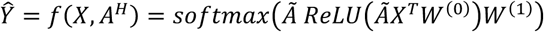

where *W*^(0)^ ∈ *R*^*m×h*^ is the input-to-hidden weight matrix projecting the *m* features input data into an *h* dimension hidden layer, *ReLU* stands for the rectified linear unit activation function, *W*^(1)^ ∈ *R*^*h×F*^ is a hidden-to-output weight matrix, and *Ŷ* ∈ *R*^*N×F*^ is the predicted probabilities of cell labels. The *softmax* activation function is defined as

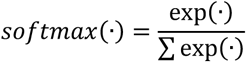

Given the reference data, we use the cross-entropy error the evaluate the predictions, *i.e*.

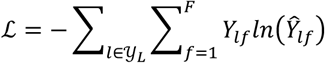

After training the model we have

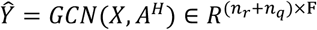

The prediction *Ŷ* is the probability of each cell within each class. The final label for cell *i* is determined as the *f*’th label when *ŷ*_*i,l*_ ≥ 0.5 where *ŷ*_*i,l*_ ∈ *Ŷ*.

When applying the scGCN model, we randomly split the reference data as training (80%), test (10%), and validation set (10%), while the query data is unlabeled that can be predicted through the above semi-supervised GCN model. For datasets in this study, we train three-layer scGCN models for a maximum of 200 epochs using Adam with a learning rate of 0.01 and early stopping with a window size of 10. For the number of hidden units, we respectively check 32, 64, 128, 256, 528,1024 and select the optimal one.

### Cross-species classification

We use the HomoloGene databases provided by NCBI (Build 68) to identify homologous genes between human and mouse, and keep only genes that have a one-to-one correspondence, which serves as a look-up table. After obtaining the intersection gene set between the reference data and the look-up table, the gene names are then converted to human gene names to obtain a compatible input for classification.

### Methods comparison

For Seurat, we identify anchors and classify cells using the “FindTransferAnchors” and “TransferData” functions respectively. We use the function “propagateLabels” in Conos to propagate labels from one annotated sample to the other unknown sample. For benchmarking with scmap, we identify the classification of query data using the “scmapCluster” function of scmap. For CHETAH, we identify the classification of query data using the “CHETAHclassifier” function. All methods are applied with default parameters. In this study, statistical significance is defined using the two-sided Wilcox test.

## Data availability

All single-cell RNA-seq datasets were downloaded from their public accessions. Specifically, the PBMC data from 6 different sequencing protocols are available from the Broad Institute Single Cell portal (https://portals.broadinstitute.org/single_cell/study/SCP424/single-cell-comparison-pbmc-data). Processed and organized data are obtained from Abdelaal T *et al* [32]. We select these single-cell data as they have been frequently used to evaluate the performance of transferring labels by different methods [27, 52-54]. The following shows the details of scRNA-seq and scATAC-seq data used in this study.

### A549 data

The A549 dataset is downloaded from GEO, which includes 3260 cells profiled using the sci-CAR protocol [55], with both scRNA-seq (GSM3271040) and scATAC-seq data (GSM3271041). After quality control, the resulted data consists of 2641 cells in our analysis.

#### Mouse brain data

The single-cell ATAC-seq dataset of adult mouse brain cells is provided by 10x Genomics, which is available through the 10x Genomics website. scRNA-seq data is obtained from the same biological system (the adult mouse brain), which can be downloaded from Allen Institute website.

#### Kidney data

The kidney dataset is downloaded from GEO, which includes 11, 296 single cells from the mammalian kidney using the sci-CAR protocol [55]. The scATAC-seq data (GSM3271045) and scRNA-seq data (GSM3271044) are profiled using the sci-CAR protocol [55]. Duplicate genes are merged by maximum value and cells that are labeled as NA are removed. Moreover, cells with less than 500 expressed genes and genes expressed in fewer than 10 cells are removed from the scRNA-seq data. Cells with less than 200 accessible loci and loci opened in fewer than 10 cells are removed from scATAC-seq data. The final dataset with 8, 837 cells is used in our analysis.

#### Lung data

The lung dataset contains lung count matrices from three RNA datasets (two lung samples from Tabula Muris, one lung sample from Han *et al* [56]), and two lung sci-ATAC-seq replicates from Cusanovich *et al* [57].

## Code availability

All the functions mentioned above are implemented as a python software, which can be downloaded at https://github.com/QSong-github/scGCN.

## Acknowledgements

The authors acknowledge the DEMON high performance computing (HPC) cluster, the Texas Advanced Computing Center (TACC) at The University of Texas at Austin (http://www.tacc.utexas.edu), and the Extreme Science and Engineering Discovery Environment (XSEDE, which is supported by National Science Foundation grant number ACI-1548562) for providing HPC resources that have contributed to the research results reported within this paper.

## Author contributions

QS, JS, and WZ developed the structure and arguments and wrote the manuscript. All the authors reviewed and approved the final manuscript.

## Materials & Correspondence

Correspondence and requests for materials should be addressed to J.S. or W.Z.

---

**Figure S1. Performance of CHETAH and scmap**

**A**. Heatmap of the accuracy matrix of each cell type identified by CHETAH based on the single cell dataset (SRP073767). **B**. Heatmap shows the accuracy matrix of each cell type identified by CHETAH based on the Cel-seq data as reference and the 10x V2 data as query set. **C**. UMAP projection of the aggregated data of GSE84133 by scmap. The first UMAP represents the aggregated data using the human data as the reference and mouse data as the query data. The second row uses the mouse data as reference and human data as query.

**Figure S2. Performance of Conos on A549 dataset**

**A**. UMAP plots of A549 cells colored by dataset (scRNAseq, scATAC-seq) and by cell states (0h, 1h, 3h), after integration by Conos. **B**. KRT7 and WDR60 are shown as overrepresented DNA motifs that are identified in 3h-specific accessibility peaks, which also exhibit overexpression in 3h cells.

**Figure S3. Application of scGCN to mouse brain dataset**

**A**. Overrepresented DNA motifs are identified in L4-specific accessibility peaks, with Foxp1, Egr3, and Smad3 motifs as the most highly enriched motifs. **B**. These motifs also exhibit upregulated expression in L4 cell subtype.

## References

1. Baron, M., et al., A Single-Cell Transcriptomic Map of the Human and Mouse Pancreas Reveals Inter-and Intra-cell Population Structure. Cell Syst, 2016. 3(4): p. 346–360 e4.

2. Puram, S.V., et al., Single-Cell Transcriptomic Analysis of Primary and Metastatic Tumor Ecosystems in Head and Neck Cancer. Cell, 2017. 171(7): p. 1611–1624 e24.

3. Athanasiadis, E.I., et al., Single-cell RNA-sequencing uncovers transcriptional states and fate decisions in haematopoiesis. Nat Commun, 2017. 8(1): p. 2045.

4. Azizi, E., et al., Single-Cell Map of Diverse Immune Phenotypes in the Breast Tumor Microenvironment. Cell, 2018. 174(5): p. 1293–1308 e36.

5. Cusanovich, D.A., et al., A Single-Cell Atlas of In Vivo Mammalian Chromatin Accessibility. Cell, 2018. 174(5): p. 1309–1324 e18.

6. Muraro, M.J., et al., A Single-Cell Transcriptome Atlas of the Human Pancreas. Cell Syst, 2016. 3(4): p. 385–394 e3.

7. Tabula Muris, C., et al., Single-cell transcriptomics of 20 mouse organs creates a Tabula Muris. Nature, 2018. 562(7727): p. 367–372.

8. Buenrostro, J.D., et al., Integrated Single-Cell Analysis Maps the Continuous Regulatory Landscape of Human Hematopoietic Differentiation. Cell, 2018. 173(6): p. 1535–1548 e16.

9. Macaulay, I.C., C.P. Ponting, and T. Voet, Single-cell multiomics: multiple measurements from single cells. Trends in Genetics, 2017. 33(2): p. 155–168.

10. Mezger, A., et al., High-throughput chromatin accessibility profiling at single-cell resolution. Nature communications, 2018. 9(1): p. 1–6.

11. Buenrostro, J.D., et al., Single-cell chromatin accessibility reveals principles of regulatory variation. Nature, 2015. 523(7561): p. 486–490.

12. Buenrostro, J.D., et al., Integrated single-cell analysis maps the continuous regulatory landscape of human hematopoietic differentiation. Cell, 2018. 173(6): p. 1535-1548. e16.

13. Satpathy, A.T., et al., Massively parallel single-cell chromatin landscapes of human immune cell development and intratumoral T cell exhaustion. Nature Biotechnology, 2019. 37(8): p. 925–936.

14. Schaum, N., et al., Single-cell transcriptomics of 20 mouse organs creates a Tabula Muris. Nature, 2018. 562(7727): p. 367–372.

15. Regev, A., et al., Science forum: the human cell atlas. Elife, 2017. 6: p. e27041.

16. Brennecke, P., et al., Accounting for technical noise in single-cell RNA-seq experiments. Nature Methods, 2013. 10(11): p. 1093–1095.

17. Lun, A.T.L. and J.C. Marioni, Overcoming confounding plate effects in differential expression analyses of single-cell RNA-seq data. Biostatistics, 2017. 18(3): p. 451–464.

18. Colomé-Tatché, M. and F.J. Theis, Statistical single cell multi-omics integration. Current Opinion in Systems Biology, 2018. 7: p. 54–59.

19. Svensson, V., Droplet scRNA-seq is not zero-inflated. Nature Biotechnology, 2020. 38(2): p. 147–150.

20. Lun, A.T., K. Bach, and J.C. Marioni, Pooling across cells to normalize single-cell RNA sequencing data with many zero counts. Genome Biol, 2016. 17: p. 75.

21. Vieth, B., et al., powsimR: power analysis for bulk and single cell RNA-seq experiments. Bioinformatics, 2017. 33(21): p. 3486–3488.

22. Grun, D., L. Kester, and A. van Oudenaarden, Validation of noise models for single-cell transcriptomics. Nat Methods, 2014. 11(6): p. 637–40.

23. Lun, A.T. and J.C. Marioni, Overcoming confounding plate effects in differential expression analyses of single-cell RNA-seq data. Biostatistics, 2017. 18(3): p. 451–464.

24. Cao, J., et al., Comprehensive single-cell transcriptional profiling of a multicellular organism. Science, 2017. 357(6352): p. 661–667.

25. Rosenberg, A.B., et al., Single-cell profiling of the developing mouse brain and spinal cord with split-pool barcoding. Science, 2018. 360(6385): p. 176–182.

26. Macosko, E.Z., et al., Highly parallel genome-wide expression profiling of individual cells using nanoliter droplets. Cell, 2015. 161(5): p. 1202–1214.

27. Butler, A., et al., Integrating single-cell transcriptomic data across different conditions, technologies, and species. Nat Biotechnol, 2018. 36(5): p. 411–420.

28. Stuart, T., et al., Comprehensive Integration of Single-Cell Data. Cell, 2019. 177(7): p. 1888-1902.e21.

29. Barkas, N., et al., Joint analysis of heterogeneous single-cell RNA-seq dataset collections. Nature Methods, 2019. 16(8): p. 695–698.

30. Kiselev, V.Y., A. Yiu, and M. Hemberg, scmap: projection of single-cell RNA-seq data across data sets. Nat Methods, 2018. 15(5): p. 359–362.

31. de Kanter, J.K., et al., CHETAH: a selective, hierarchical cell type identification method for single-cell RNA sequencing. Nucleic Acids Research, 2019. 47(16): p. e95–e95.

32. Abdelaal, T., et al., A comparison of automatic cell identification methods for single-cell RNA sequencing data. Genome biology, 2019. 20(1): p. 194.

33. Kipf, T. and M. Welling, Semi-supervised classification with graph convolutional networks ICLR. 2017.

34. Yuan, Y. and Z. Bar-Joseph, GCNG: Graph convolutional networks for inferring cell-cell interactions. bioRxiv, 2019: p. 2019.12.23.887133.

35. Li, J., et al., Neural inductive matrix completion with graph convolutional networks for miRNA-disease association prediction. Bioinformatics, 2020. 36(8): p. 2538–2546.

36. Zeng, Y., et al., Accurately Clustering Single-cell RNA-seq data by Capturing Structural Relations between Cells through Graph Convolutional Network. bioRxiv, 2020: p. 2020.09.02.278804.

37. Zhao, T., et al., Identifying drug–target interactions based on graph convolutional network and deep neural network. Briefings in Bioinformatics, 2020.

38. Becht, E., et al., Dimensionality reduction for visualizing single-cell data using UMAP. Nature biotechnology, 2019. 37(1): p. 38–44.

39. Haghverdi, L., et al., Batch effects in single-cell RNA-sequencing data are corrected by matching mutual nearest neighbors. Nature biotechnology, 2018. 36(5): p. 421–427.

40. Rousseeuw, P.J., Silhouettes: A graphical aid to the interpretation and validation of cluster analysis. Journal of Computational and Applied Mathematics, 1987. 20: p. 53–65.

41. Lützner, N., et al., FOXO3 is a glucocorticoid receptor target and regulates LKB1 and its own expression based on cellular AMP levels via a positive autoregulatory loop. PloS one, 2012. 7(7): p. e42166.

42. Mostafa, M.M., et al., Glucocorticoid-driven transcriptomes in human airway epithelial cells: commonalities, differences and functional insight from cell lines and primary cells. BMC medical genomics, 2019. 12(1): p. 1–21.

43. Bhadri, V.A., et al., Evaluation of the NOD/SCID xenograft model for glucocorticoid-regulated gene expression in childhood B-cell precursor acute lymphoblastic leukemia. BMC Genomics, 2011. 12(1): p. 565.

44. Caldwell, J.M., et al., Glucocorticoid-regulated genes in eosinophilic esophagitis: a role for FKBP51. The Journal of allergy and clinical immunology, 2010. 125(4): p. 879-888.e8.

45. Song, Q., et al., Dissecting intratumoral myeloid cell plasticity by single cell RNA-seq. Cancer medicine, 2019. 8(6): p. 3072–3085.

46. Song, Q., et al., scLM: automatic detection of consensus gene clusters across multiple single-cell datasets. bioRxiv, 2020: p. 2020.04.22.055822.

47. Ghosh, A. and D. Kandasamy, Interpretable Artificial Intelligence: Why and When. AJR Am J Roentgenol, 2020. 214(5): p. 1137–1138.

48. Moore, J.H., et al., Preparing next-generation scientists for biomedical big data: artificial intelligence approaches. Per Med, 2019. 16(3): p. 247–257.

49. Filipp, F.V., Opportunities for Artificial Intelligence in Advancing Precision Medicine. Curr Genet Med Rep, 2019. 7(4): p. 208–213.

50. Haghverdi, L., et al., Batch effects in single-cell RNA-sequencing data are corrected by matching mutual nearest neighbors. Nat Biotechnol, 2018. 36(5): p. 421–427.

51. Dudoit, S., J. Fridlyand, and T.P. Speed, Comparison of Discrimination Methods for the Classification of Tumors Using Gene Expression Data. Journal of the American Statistical Association, 2002. 97(457): p. 77–87.

52. Kiselev, V.Y., A. Yiu, and M. Hemberg, scmap: projection of single-cell RNA-seq data across data sets. Nature Methods, 2018. 15(5): p. 359–362.

53. Stuart, T., et al., Comprehensive integration of single-cell data. Cell, 2019. 177(7): p. 1888-1902. e21.

54. Abdelaal, T., et al., A comparison of automatic cell identification methods for single-cell RNA sequencing data. Genome Biology, 2019. 20(1): p. 194.

55. Cao, J., et al., Joint profiling of chromatin accessibility and gene expression in thousands of single cells. Science, 2018. 361(6409): p. 1380–1385.

56. Han, X., et al., Mapping the Mouse Cell Atlas by Microwell-Seq. Cell, 2018. 172(5): p. 1091-1107.e17.

57. Cusanovich, D.A., et al., Multiplex single-cell profiling of chromatin accessibility by combinatorial cellular indexing. Science, 2015. 348(6237): p. 910–914.

