## Supplementary figures and images for "scGCN: a Graph Convolutional Networks Algorithm for Knowledge Transfer in Single Cell Omics"

### Supplement Figures

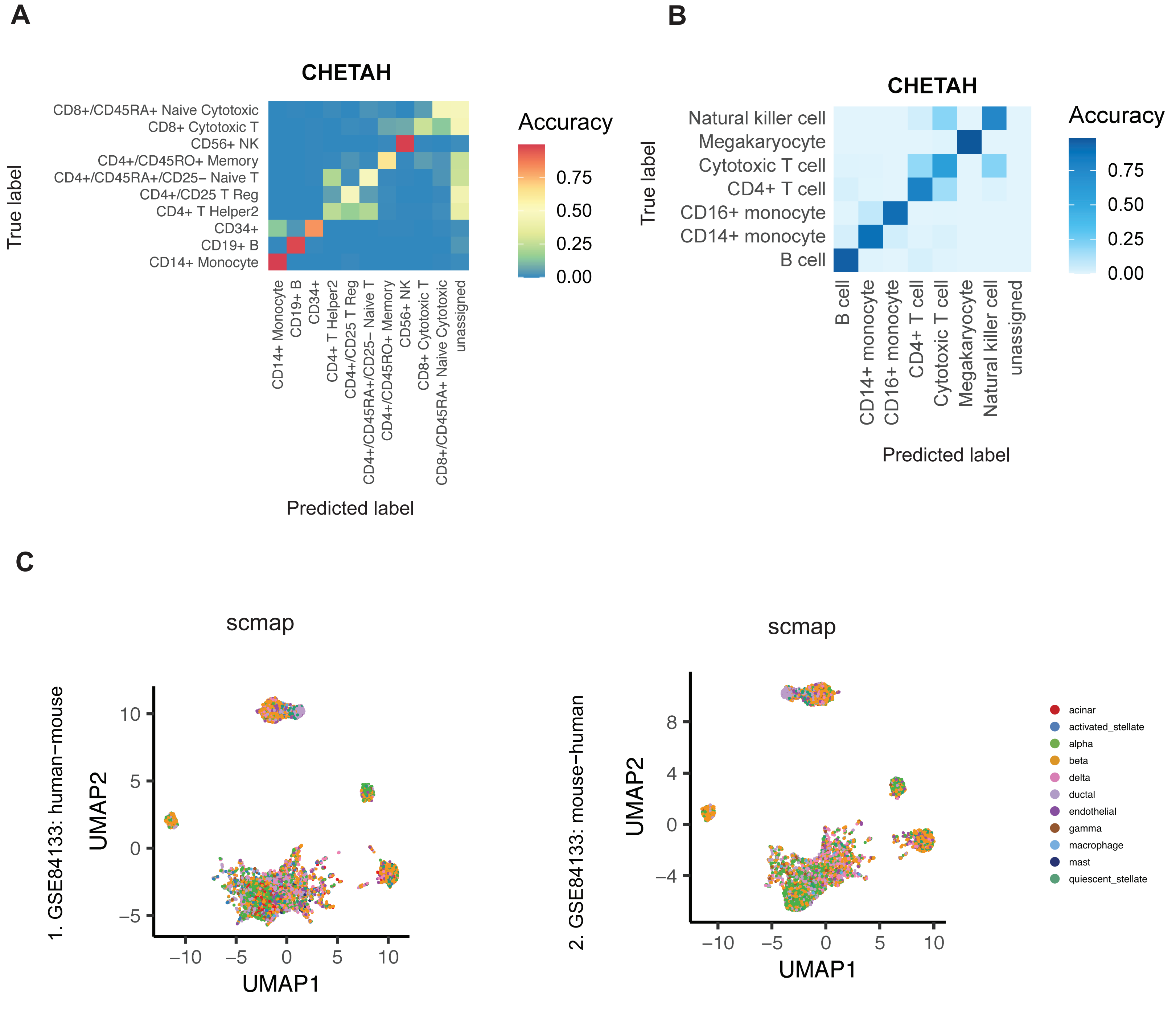

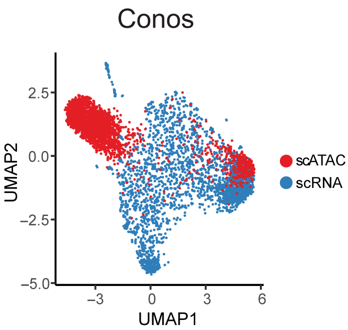

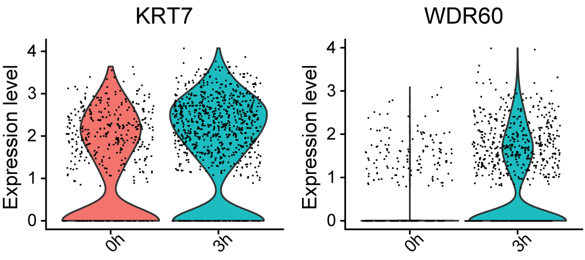

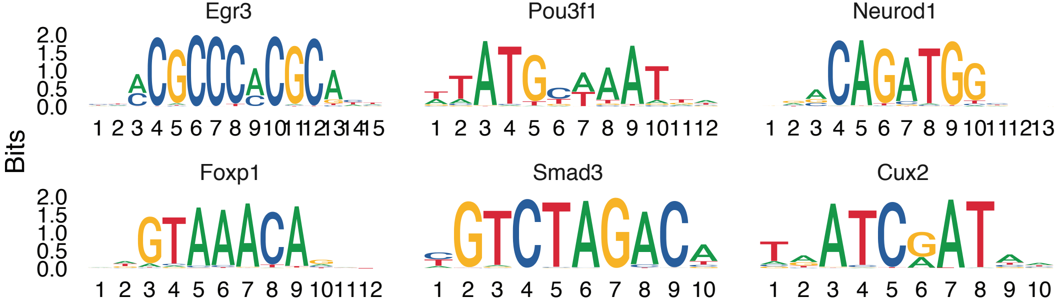

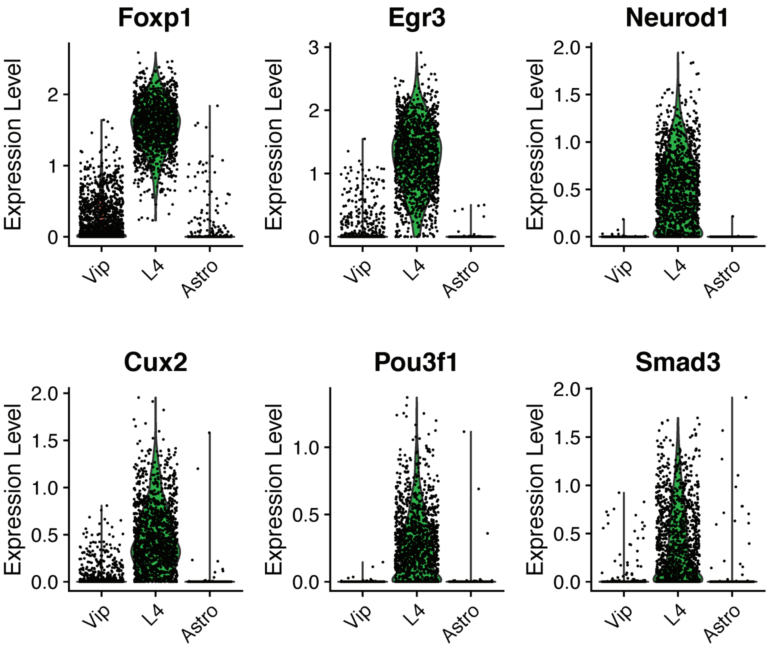
